# Interactions with a phage gene underlie costs of a β-lactamase

**DOI:** 10.1101/2023.09.23.559150

**Authors:** Huei-Yi Lai, Tim F. Cooper

## Abstract

The fitness cost of an antibiotic resistance gene (ARG) can differ across host strains creating refuges that allow maintenance of an ARG in the absence of direct selection for its resistance phenotype. Despite the importance of such ARG-host interactions for predicting ARG dynamics, the basis of ARG fitness costs and their variability between hosts are not well understood. We determined the genetic basis of a host-dependent cost of a β-lactamase, *bla*_TEM-116*_, that conferred a significant cost in one *Escherichia coli* strain but was close to neutral in 11 other *Escherichia spp.* strains. Selection of a *bla*_TEM-116*\**_ encoding plasmid in the strain in which it initially had a high cost resulted in rapid and parallel compensation to that cost through mutations in a P1 phage gene, *relA_P1_*. When the wildtype *relA_P1_* gene was added to a strain in which it was not present and in which *bla*_TEM-116*\**_ was neutral, it caused the ARG to become costly. Thus, *relA_P1_* is both necessary and sufficient to explain *bla*_TEM-116*\**_ costs in at least some host backgrounds. To our knowledge, these findings represent the first demonstrated case of the cost of an ARG being influenced by a genetic interaction with a phage gene. The interaction between a phage gene and a plasmid-borne ARG highlights the complexity of selective forces determining the maintenance and spread of ARGs, and, by extension, encoding phage and plasmids, in natural bacterial communities.

## Introduction

The spread of antibiotic resistance in pathogenic bacteria is a growing problem to public health (Collaborators et al. 2022). Although increases in resistance traits is mainly attributed to selection imposed by exposure to corresponding antibiotics, other mechanisms may also contribute to the success of antibiotic resistance traits (Björkman and Andersson 2000; Gelder et al. 2004; Rahman et al. 2023). For example, in the absence of antibiotic selection, the genetic determinants of antibiotic resistance, antibiotic resistance mutations (ARMs) and antibiotic resistance genes (ARGs), can impose a fitness cost on bacteria (Andersson and Hughes 2010). These costs vary between strains, resulting in a relative advantage for some resistant strains in the absence of antibiotic selection (Björkman and Andersson 2000; Andersson and Hughes 2010). It is also possible that costs can be compensated, perhaps at different rates and extents in different strains (Andersson and Hughes 2010; Barrick et al. 2010). Understanding the costs of ARGs, and their potential to change, is crucial to understanding, and eventually predicting, the ecological and evolutionary dynamics of resistance traits in bacterial populations.

Whereas ARMs generally occur in host cell genes, ARGs are disseminated widely across bacterial species so that genetic interactions that determine their fitness cost will be diverse and can have a significant influence on their selection (Moradigaravand et al. 2018). Candidate factors linked to the success of horizontally acquired ARGs include: compatibility of GC content and codon usage with the host (Popa et al. 2011; Moradigaravand et al. 2018), gene function in the context of the host’s metabolism (Rivera et al. 1998), protein interactivity (Jain et al. 1999), toxicity (Sorek et al. 2007), gene expression level (Park and Zhang 2012), and the 5’ UTR stability of mRNA (Kudla et al. 2009; Goodman et al. 2013). The effect of these factors on the costs of ∼200 clinically prevalent ARGs has been examined in *Escherichia coli* (Porse et al. 2018). That study found costs could sometimes be predicted. For example, the fitness cost of cell-interacting ARGs, such as efflux proteins, were positively correlated with the phylogenetic distance between the donor bacterium and *E. coli* (Porse et al. 2018). However, no such correlation was found for drug-interacting ARGs, such as drug modification enzymes, underlining the complexity of mechanisms likely to influence ARG selection in natural populations (Porse et al. 2018). Moreover, to our knowledge, no general mechanisms have been proposed to explain differences in the cost of a single ARG over different host strains of the same bacterial species.

Detailed studies on individual ARGs have revealed diverse specific mechanisms underlying fitness costs. For example, the cost of the *tetAR* tetracycline resistance operon has been attributed to its unregulated expression in the absence of tetracycline (Nguyen et al. 1989; Rajer and Sandegren 2022). Costs of the β-lactamases BlaCTX-M-15, BlaSME-1, BlaVIM-1 and BlaSPM-1 are in part due to their signal peptides, which probably decrease host fitness by disrupting their cell membrane (Marciano et al. 2007; López et al. 2019; Rajer and Sandegren 2022). By contrast, expression of the β-lactamases BlaOXA and BlaSFO-1 is associated with changes in host cell peptidoglycan composition, suggesting the residual DD-transpeptidase activity of β-lactamases may underlie their costs (Fernández et al. 2012).

Whatever the basis of the cost of a particular ARG, those costs are themselves subject to evolutionary change. Mutations that compensate for the cost of ARM-mediated resistance occur repeatedly in laboratory evolution experiments and have been identified in some clinical isolates (Comas et al. 2012; Harrison et al. 2015; Millan et al. 2015; Loftie-Eaton et al. 2017; Lin et al. 2018; Merker et al. 2018). These compensatory mutations often occur in genes encoding products that physically interact with those of the focal ARG, implying that studying the basis of compensation to an ARG’s cost may help to predict mechanisms through which that cost occurs (Davis et al. 2009; Szamecz et al. 2014). For example, studies on different types of ARG-carrying plasmids have found compensatory mutations that effect transcription and DNA structure, suggesting that these processes are a common cause of plasmid cost (Comas et al. 2012; Millan et al. 2015; Loftie-Eaton et al. 2017; Merker et al. 2018; Wein et al. 2019).

Previously we found the β-lactamase gene, *bla*_TEM-116*\**_, imposed a significant fitness cost of greater than 10% in the *E. coli* strain M114 but was only slightly costly or nearly neutral in 11 other *Escherichia spp.* host strains (Lai and Cooper 2023). Here we examine the potential for compensation of this cost. We find that replicate M114 populations containing a *bla*_TEM-116*\**_ plasmid evolved to reduce rates of population-level plasmid loss over time. This change was due to compensatory mutations acting to reduce the fitness cost of *bla*_TEM-116*\**_. Genome sequencing and genetic reconstruction experiments demonstrated that compensation was due to mutations effecting a *relA* orthologue encoded by a phage-like P1 element and that this gene was sufficient to cause *bla*_TEM-116*\**_ to confer a cost in a strain in which it was initially neutral.

## Materials and Methods

### Bacterial strains, plasmids and media

*E. coli* strains M114 and REL606 have been described previously (Wang et al. 2016). Strains were propagated in Davis Mingioli medium supplemented with 250 mg/ml glucose (DM250) at 37°C for all selection and fitness estimation experiments unless noted otherwise. Lysogeny broth (LB) supplemented with ampicillin (Ap; 100 μg/ml) as appropriate to ensure plasmid selection was used for growth as part of genetic modification protocols. Media used to grow strains containing the plasmids pmFP and pmFP-*bla*_TEM-116_*_*_* was supplemented with 50 mg/ml kanamycin as described in the text.

Construction and features of pmFP and pmFP-*bla*_TEM-116*\**_ have been described previously (Lai and Cooper 2023). Briefly, pmFP is a derivative of pUA66 from which the promoterless *gfp* has been removed (Zaslaver et al. 2006; Jiang et al. 2015). The *bla*_TEM-116*\**_ gene and promoter was cloned into pmFP from pTarget to produce pmFP-*bla*_TEM-116*\**_ (Zaslaver et al. 2006; Jiang et al. 2015). Sequencing of this constructfound that the *bla*_TEM-116_ gene present in pTarget differed from the reference sequence (NCBI accession U36911) by a non-synonymous mutation (Q274R) and we add an asterisk to the genotype of the allele used here to indicate this.

### Selection for ARG compensation

To select for compensation of plasmid costs we inoculated replicate cultures with colonies of relevant strains obtained by streaking from freezer stocks. Initial cultures were grown with antibiotics as appropriate to ensure plasmids were present in all cells. Cultures were propagated by daily 1:100 dilution into fresh antibiotic-free media for 61 cycles (∼400 generations). At the end of this time, cultures were inoculated into DM250 supplemented with kanamycin to kill any plasmid-free cells, thereby resetting plasmid-containing cells to fixation. These ‘reset’ cultures were then inoculated into new antibiotic-free DM250 media and propagated by daily 1:100 dilution into fresh antibiotic-free media for a further 21 cycles (∼140 generations). Plasmid frequency was determined periodically throughout the second part of the experiment and compared to control populations started with the same plasmid-containing host strains taken directly from freezer stocks (i.e., omitting the 61-cycle initial evolution treatment). Plasmid frequency was determined by plating population samples on selective and non-selective media to determine plasmid-containing cell and total cell counts, respectively.

### Introducing mutations into relA_P1_

Genetic modification of the P1-like phage gene, *relA_P1_*, was performed using the portMAGE protocol (Nyerges et al. 2016). Briefly, the portMAGE2 plasmid was used to transform target cells. Transformants were maintained at 30°C to prevent induction of the encoded *c*I857-repressed λ red recombinase enzymes. Overnight cultures of transformants were diluted 1:200 into 5 ml LB supplemented with Ap and grown for three hours. To induce red recombination enzymes and a transitive mutator phenotype, these cells were induced at 42 °C for 15 minutes before being placed on ice for 10 minutes and then made electrocompetent by washing three times with 10 ml ice-cold Milli-Q water. Washed cells were resuspended in 100 μL Milli-Q water and placed on ice before mixing with an oligonucleotide (1 μM final concentration) designed to introduce focal mutations into the target cell. Oligonucleotides were designed following published guidelines and are listed in Table S1 (Gallagher et al. 2014). After electroporation, cells were recovered at 30°C in LB + Ap for 2 hours before either plating on LB + Ap agar plates or inoculating into 5 mL LB + Ap for use in a subsequent portMAGE cycle. Four cycles of portMAGE were performed before genotyping randomly chosen clones to test for target modifications. To cure the portMAGE2 plasmid, cells were cultured at 42°C overnight on LB agar, then restreaked to fresh LB agar and incubated at 37°C before screening to identify ampicillin susceptible strains.

### relA_P1_ integration at attTn7 site

To integrate *relA_P1_* into REL606, the *relA_P1_* gene, including flanking sequences extending 134 bp upstream and 140 bp downstream of the open reading frame, was cloned into the mini-Tn*7* delivery vector pGRG36 (McKenzie and Craig 2006). The resulting plasmid, pGRG36-*relA_P1_*, was used to transform REL606 and transformants were selected on LB + Ap agar at 30°C. To induce mini-Tn*7* transposition, transformants were inoculated into LB broth supplemented with 0.1% arabinose and grown overnight at 30°C. To cure cells of pGRG36, colonies were grown and restreaked on LB agar. Integration of *relA_P1_* at the *att*Tn*7* site was confirmed by PCR.

### Fitness competitions

Flow cytometry-based competition assays were used to estimate the fitness cost of a plasmid or a gene following a protocol described previously (Lai and Cooper 2023). Briefly, cells were inoculated from freezer stocks into DM250 supplemented with kanamycin if they carried a plasmid and incubated overnight. These cultures were diluted 1:100 into fresh antibiotic-free DM250 for two 24-hour pre-conditioning cycles. After pre-conditioning, equal volumes of competing test and reference pUA66-*P_rpsL_gfp* plasmid-containing cells were mixed and competitor proportions determined using flow cytometry (day 0). The cell mixture was also diluted 1:100 into fresh medium and cultured overnight before again determining the proportion of both competitors (day 1). Estimates of the fitness effect of focal plasmids or ARGs were made indirectly as the difference in fitness of competitors differing by that focal genetic trait competed separately against a common pUA66-*P_rpsL_gfp* plasmid-containing reference strain. For example, the difference in fitness of a strain containing the empty pmFP vector relative to the reference strain, and the fitness of the same strain containing the pmFP*-bla*_TEM-116*\**_ plasmid relative to the reference strain, was used to isolate the fitness effect of the *bla*_TEM-116*\**_ ARG.

### Whole genome sequencing of evolved clones

One plasmid-carrying clone was isolated from each M114 evolved line. Genomic DNA of the evolved clones was isolated using the Wizard® Genomic DNA Purification Kit (Promega) and short-read sequenced using the Illumina HiSeq platform (Microbial Genomic Sequencing Center, USA). Breseq was used to identify mutations relative to the ancestral strain (Deatherage and Barrick 2014).

### Statistics

The R statistical computing platform version 4.3.1 was used for all analysis and visualization (R Core Team 2021). The flowCore package was used to analyze flow cytometry data and the Binom.confit function from the binom package was used to calculate confidence intervals on proportions (Ellis et al. 2022).

## Results

Variation in fitness costs leads to different ecological and evolutionary dynamics of the bla_TEM-116*_ plasmid Variation in the fitness cost of a given ARG across different bacterial strains is expected to affect the strength of both purifying selection and selection for compensation of costs. As a first step to test this expectation, we examined the stability of a non-conjugative plasmid, pmFP, encoding *bla*_TEM-116*\**_ in two hosts, REL606 and M114, in which it confers different costs (cost in REL606: 0.3% (±1.0 CI95); cost in M114: 10.9% (±3.3 CI95)).

Replicate lines started with M114 carrying either pmFP-*bla*_TEM-116*\**_ or a pmFP vector-only control plasmid were initially propagated for 400 generations without any direct selection for the plasmids. In the M114 populations, in which the pmFP-*bla*_TEM-116*\**_ plasmid had a high cost, it remained present in an average of only 4.5% (0.2 – 11.1 CI95) of cells at the end of the selection period. By contrast, the control plasmid remained present in 93.9% of cells (86.8 – 97.5 CI95). The small number of cells retaining the pmFP-*bla*_TEM-116*\**_ plasmid is consistent with strong purifying selection against plasmid carriage and, therefore, the potential for selection of compensatory mutations that reduce the cost of carriage.

To test if selected lines evolved in a way that could influence ongoing plasmid dynamics, for example through selection of compensatory mutations that reduce purifying selection against plasmid-containing cells, we exposed them to kanamycin to select for remaining plasmid-carrying cells in each line. These populations, in which plasmid-containing cells were reset to fixation, were used to start new selection lines that were propagated through an additional 21 daily transfers. Loss of pmFP-*bla*_TEM-116*\**_ during this second selection period was greatly reduced in M114 lines that had been ‘pre-evolved’ for 400 generations compared to naïve control lines in which no previous evolution had occurred (pre-evolved: plasmid containing cells at 21 days = 93.7%, 86.6 – 97.4 CI95; naïve: 7.9%, 3.7 – 15.5 CI95; Fig. 1). In lines started with the host strain REL606, in which *bla*_TEM-116*\**_ conferred no detectable cost, a period of pre-evolution had no effect on plasmid carriage (pre-evolved: plasmid containing cells at 21 days = 60.9%, 50.6 – 70.3 CI95; naïve: 66%, 55.8 – 75.0 CI95; Fig. 1). These results indicate that some evolutionary change occurred during the initial evolution of M114(pmFP-*bla*_TEM-116*\**_) lines that increased population-level maintenance of the plasmid in the absence of direct selection.

**Figure 1.**
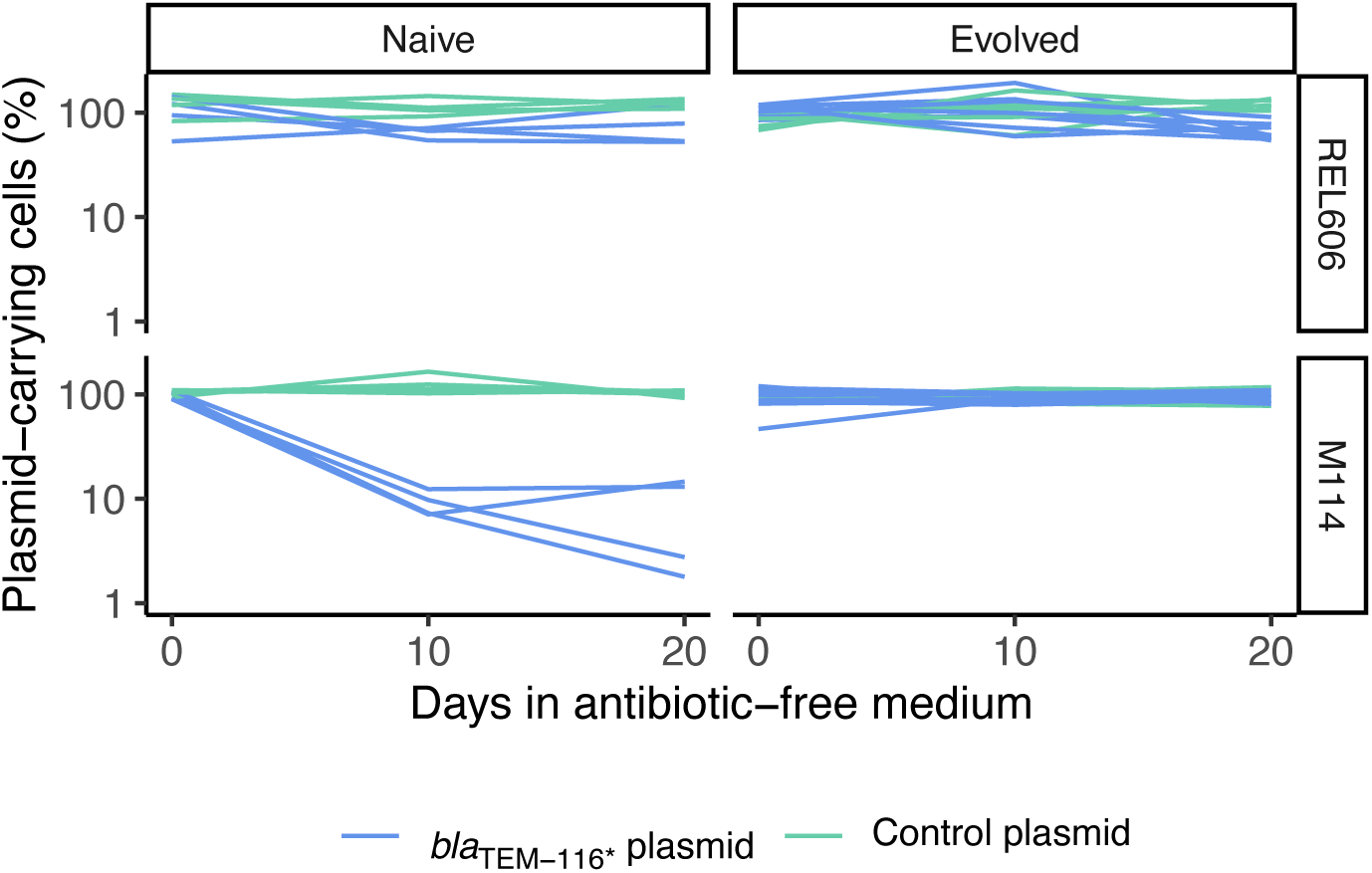
Dynamics of control and *bla_TEM-116*_* plasmids in naïve and pre-evolved populations. Dynamics of control (empty vector, pmFP) and pmFP-*bla*_TEM-116*\**_ plasmid-carrying cells was tracked in REL606 and M114 host strains. Populations were started directly from freezer stocks (left panels; naive) or after an initial period of evolution during which mutations that increased fitness of plasmid-carrying cells were expected to be selected (right panels; evolved). Naive populations: n = 4; evolved populations: n = 6 (control plasmid) or 9 (*bla*_TEM-116*\**_ plasmid).

### Compensatory mutations reduced the cost of bla_TEM-116*_ in M114

To determine if changes in pmFP-*bla*_TEM-116*\**_ dynamics in the pre-evolved M114 lines was due to compensation of its cost, we isolated a plasmid-containing clone from each line and estimated the fitness effect of the plasmid. We found the plasmid did not impose a significant fitness effect in any evolved clone, consistent with the evolution of compensating mutations (Fig. 2A). Compensatory mutations could occur in the bacterial genome or on the pmFP-*bla*_TEM-116*\**_ plasmid. To distinguish between these possibilities, we cured the evolved plasmid from the isolated clones and introduced into cured clones the ancestral pmFP-*bla*_TEM-116*\**_ plasmid. In no case did this plasmid confer a fitness cost, indicating that compensatory mutations reside in the bacterial chromosome (Fig. 2B). Further, the fitness effects of the *bla*_TEM-116*\**_ ARG by itself (given by the difference in fitness of strains carrying pmFP-*bla*_TEM-116*\**_ or the empty pmFP vector) was not different from those of the entire pmFP-*bla*_TEM-116*\**_ plasmid, indicating compensation was to the ARG itself, not some other aspect of the pmFP plasmid backbone (Fig. 2B; *t*-test, P>0.05 for all paired comparisons).

**Figure 2.**
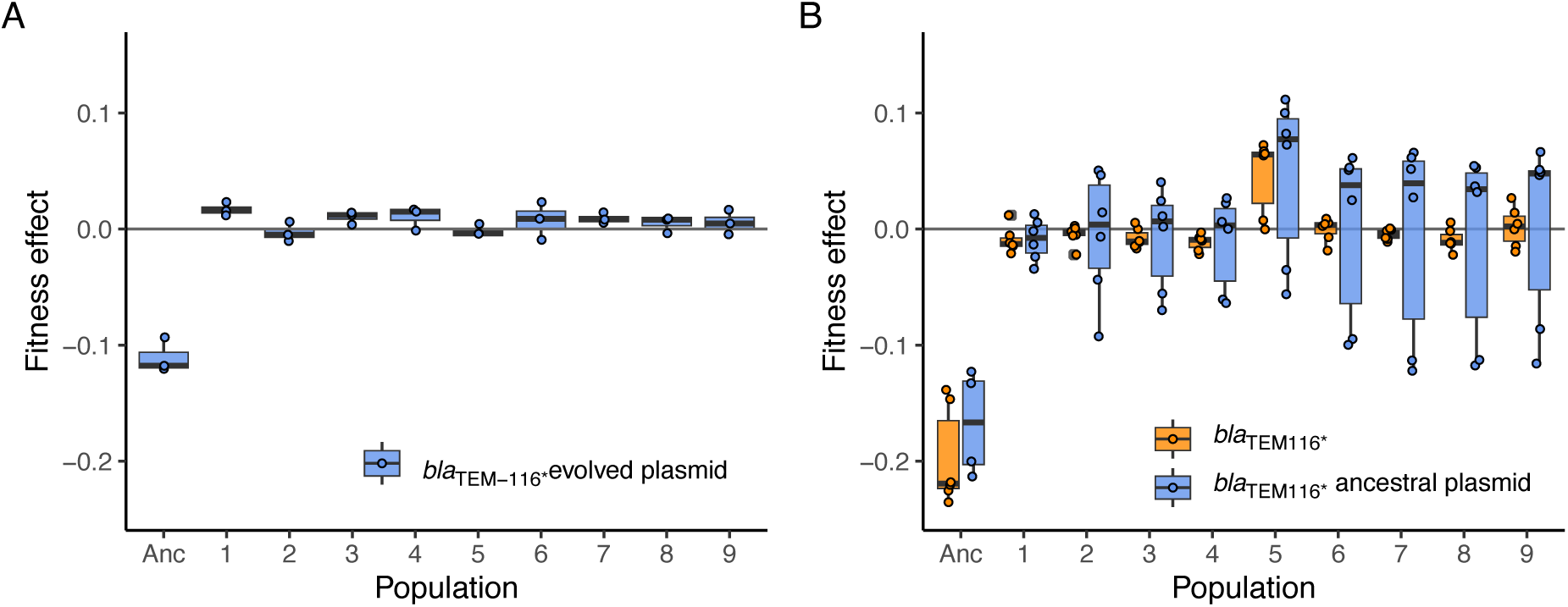
Compensation of fitness cost of *bla_TEM-116*_* in evolved M114 lines. **(A)** Fitness effect of the evolved *bla*_TEM-116*\**_ plasmid on fitness of the ancestor (Anc) and clones isolated from nine independently evolved lines. Plasmid effect was estimated from the fitness difference of isolated clones and paired derivatives that were cured of the plasmid. **(B)** Fitness effect of the ancestral *bla*_TEM-116*\**_ plasmid and the *bla*_TEM-116*\**_ ARG in the ancestral strain (Anc) and in the same independently evolved clones shown in panel A. Plasmid effect was determined by introducing the ancestral *bla*_TEM-116*\**_ plasmid into evolved clones that had been cured of the evolved plasmid. The *bla*_TEM-116*\**_ ARG effect was estimated as the difference in fitness of evolved clones carrying the ancestral *bla*_TEM-116*\**_ plasmid and paired clones carrying an empty vector. Boxes indicate mean and first and third quartiles, and symbols indicate individual estimates (A, n = 3; B, n ≥ 4).

### Mutations on a native phage alleviate the cost of the bla_TEM-116*_ plasmid

To identify the genetic basis of compensatory mutations occurring in the M114 evolved clones, we sequenced the genomes of the same pmFP-*bla*_TEM-116*\**_ plasmid-carrying clones in which we measured plasmid and ARG fitness effects (Figs. 2A & B). We found mutations involving a native 90 kb P1-like phage (hereafter, P1_M114_) in all clones. Six of the evolved *bla*_TEM-116*\**_ plasmid clones lost the entire phage while the other three clones gained a mutation, I179S, in a phage-encoded *relA* gene, a predicted ppGpp synthetase (Table S2). (To distinguish the phage-encoded *relA* from the chromosomal *E. coli relA* gene, we hereafter refer to it as *relA_P1_*.) By contrast, none of the six evolved control pmFP plasmid-containing lines had any mutations in P1_M114_ (Table S2). The enrichment of P1_M114_ mutations in *bla*_TEM-116*\**_ expressing clones makes these changes a good candidate for underlying the parallel evolved compensation trait (Figs. 2A & B).

To test if evolved P1_M114_ mutations are sufficient to compensate for the cost of pmFP-*bla*_TEM-116*\**_, we determined the plasmids cost in a derivative of the ancestral M114 strain containing the evolved *relA_P1_* I179S mutation (Fig. 3A). In this strain, the cost of the *bla*_TEM-116*\**_ plasmid was reduced from 16.1% (±0.6 CI95) in the ancestor to being neutral (0.0% ±1.0 CI95). We were unable to cure P1_M114_ from M114 cells, so we introduced a null *relA_P1_* allele into the ancestral clone to determine the effect of losing only that gene. Again, the effect of the *bla*_TEM-116*\**_ plasmid was changed to become neutral (0.0% ±1.4 CI95; Fig. 3A).

**Figure 3.**
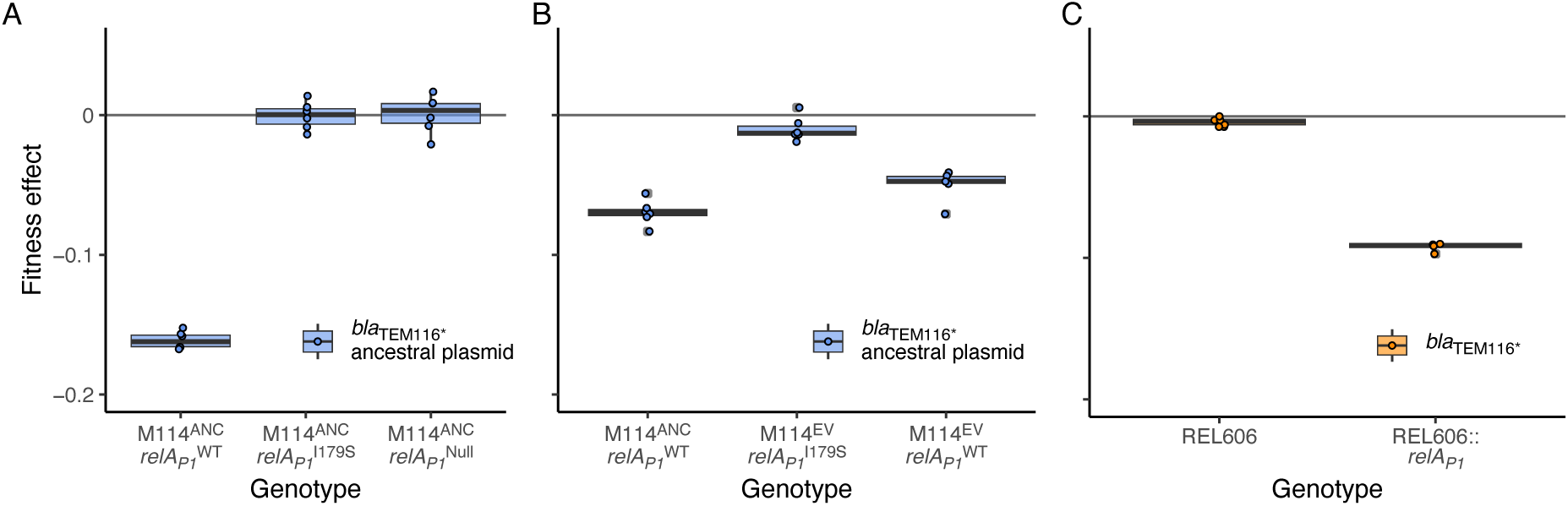
Effect of *relA_P1_* mutations on fitness effect of the *bla*_TEM-116*\**_ plasmid and ARG. **(A)** Fitness effect of the pmFP-*bla*_TEM-116*\**_ plasmid in ancestral M114 with *relA_P1_* I179S and *relA_P1_* null alleles. **(B)** Fitness effect of the pmFP-*bla*_TEM-116*\**_ plasmid in the ancestral M114 strain and derivatives of an evolved M114 clone containing either the evolved *relA_P1_* I179S allele or a reverted *relA_P1_* wild type allele. **(C)** Fitness effect of the *bla*_TEM-116*\**_ ARG in REL606 and a derivative encoding the ancestral *relA_P1_* gene transferred from M114. Boxes indicate mean and first and third quartiles, and symbols indicate individual estimates (n ≥ 5).

To further characterize the genetic basis of compensation we reverted the evolved *relA ^I179S^*allele to the ancestral allele in one evolved clone. We found that this change caused the cost of pmFP-*bla*_TEM-116*\**_ to increase significantly (difference in mean cost 4%, *t-*test P < 0.001), but not to a level as great as that seen in the ancestral strain (difference in mean cost 1.9%, *t-* test P = 0.02) (Fig. 3B). (We note evidence of a block effect such that the effect of pmFP-*bla*_TEM-116*\**_ in the ancestral M114 was less in this experiment than in others reported in this study. However, competitions reported in figure 3B were performed in the same experimental block so that they are comparable to one another.) Together our results indicate that evolved changes in *relA_P1_* are sufficient to remove all cost of pmFP-*bla_TEM-116*_* in ancestral M114 cells but that other evolved changes also contribute some component of reducing pmFP-*bla_TEM-116*_* cost in the focal evolved line.

Finally, we determined the fitness effect of *relA_P1_* alleles independent of pmFP-*bla*_TEM-116*\**_. We found that the I179S and null *relA_P1_* alleles had no effect on ancestral M114 fitness and that reverting the evolved *relA_P1_* I179S allele to the ancestral allele had no effect on fitness of an evolved strain cured of the plasmid (*t-*test all P>0.05; Fig. 4). Evidently changes to *relA_P1_* affect fitness only through a positive epistatic interaction that compensates for the cost of the *bla*_TEM-116*\**_ ARG and do not confer any general benefit to cells in the test environment.

**Figure 4.**
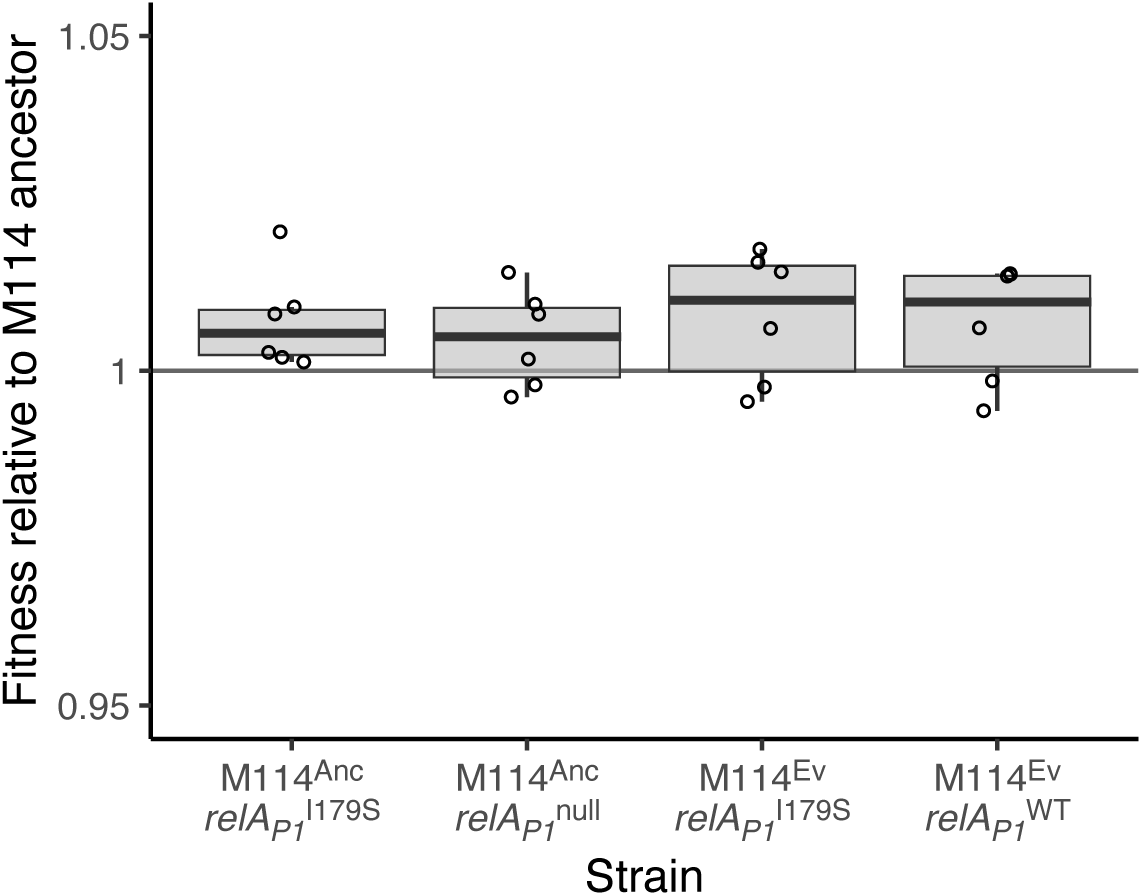
Fitness effect of *relA_P1_* in M114. Fitness of the M114 ancestor and an evolved strain with indicated *relA1_P1_* alleles relative to the M114 ancestor. In no case was a significant difference in fitness detected (*t*-test all P>0.05). Boxes indicate mean and first and third quartiles, and symbols indicate individual estimates (n ≥ 6).

**Figure 5.**
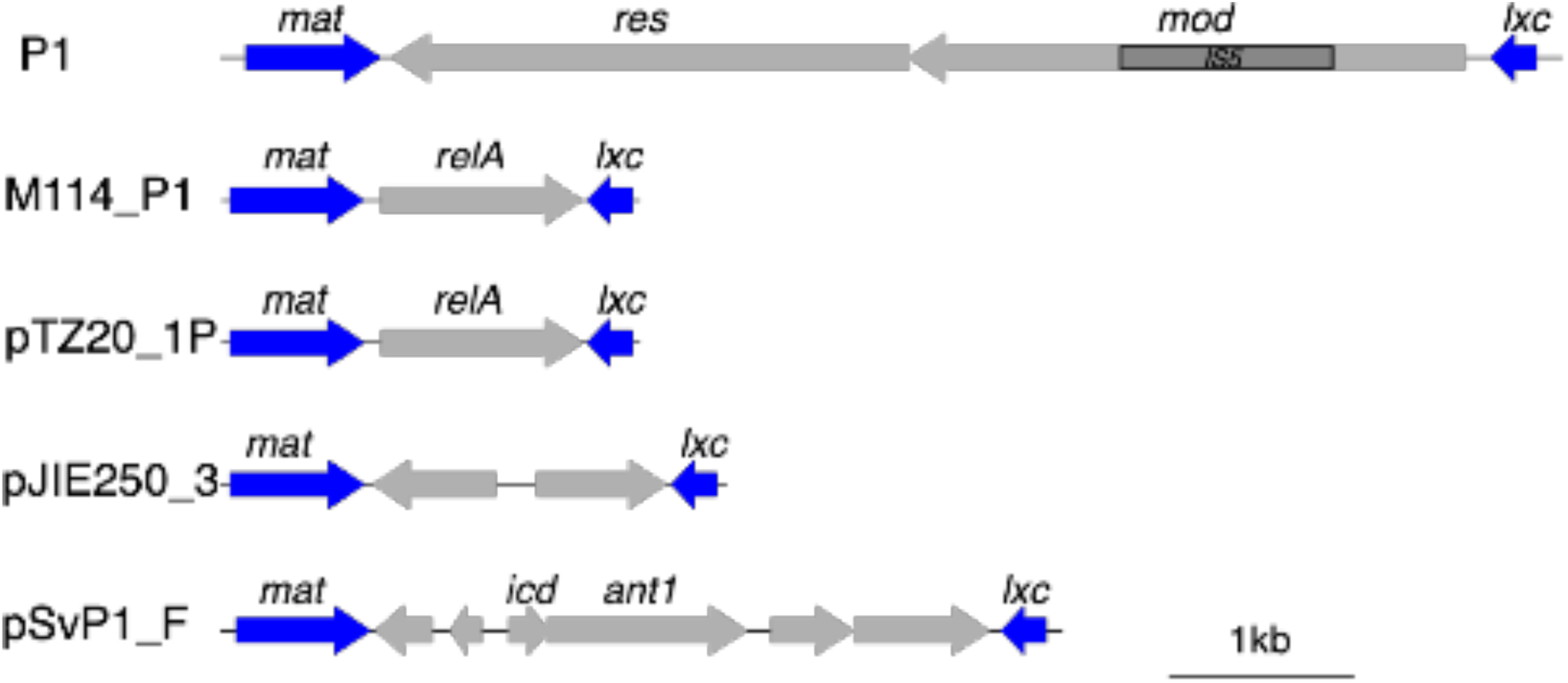
Synteny of the P1 RD-1 region shows a variable gene repertoire. A close-up of the RD-1 of phage P1 (from Fig S2). The RD-1 region is consistently flanked by *mat* and *lxc* genes, but internal gene content is variable (Dedrick et al. 2017).

### Expression of relA_P1_ can cause bla_TEM-116*_ plasmid costs

Our results demonstrate that the wild type *relA_P1_* gene interacts with the *bla*_TEM-116*\**_ gene to cause a cost in M114. To test if the same interaction can cause *bla*_TEM-116*\**_ to be costly in other genetic backgrounds we introduced *relA_P1_* into the lab strain REL606, which does not naturally have the phage P1_M114_ or *relA_P1_* and has no cost when carrying the *bla*_TEM-116*\**_ plasmid (Lai and Cooper 2023). If *relA_P1_* is sufficient to cause the *bla*_TEM-116*\**_ plasmid to confer a cost to host cells, we expected that expressing *relA_P1_* alone would incur a cost of the plasmid similar to that in M114. By contrast, if other phage P1_M114_-encoded genes or M114-specific genes are involved, we expected that expressing *relA_P1_* alone in REL606 would have little or no influence on the fitness effect of *bla*_TEM-116*\**_.

We integrated *relA_P1_* at the *att*Tn*7* site in the REL606 chromosome and measured the effect of this insertion on the cost of *bla_TEM-116*_*. We found that *bla_TEM-116*_* confers a 9% (±0.5 % 95CI, *t*-test P < 0.001) fitness cost in REL606 Tn*7*:: *relA_P1_* but has no cost in the wild type REL606 (cost = 0.2% ±0.4 % 95CI, *t*-test P = 0.08) (Fig. 3C). In the absence of any plasmid or ARG, the *relA_P1_* insertion conferred no cost in the competition environment (*t*-test P = 0.16). Taken together, the compensated cost of *bla_TEM-116*_* in M114 with a *relA_P1_* null mutant and the emergent cost of *bla_TEM-116*_* in REL606 Tn*7*:: *relA_P1_* indicates that *relA_P1_* can be necessary and sufficient to cause *bla_TEM-116*_* to confer a fitness cost.

### Similarity of relA_P1_ to chromosomal relA/spoT and prevalence in phage

RelA_P1_ shares sequence similarity to the N-terminal enzymatic domains of the *E. coli* ppGpp synthetase RelA and ppGpp synthetase/hydrolase SpoT, but lacks their C-terminal regulatory domains (Fig. S1). Sequence alignment of RelA_P1_ to RelA and SpoT showed that the mutated I179 residue of RelA_P1_ is located near residues critical for ppGpp synthetase activity, suggesting both that RelA_P1_ has functional ppGpp synthetase activity and that this activity is affected by the evolved mutation (Fig. S1). We note that the null and I179S *relA_P1_* alleles caused a similar reduction in cost of the *bla*_TEM-116*\**_ plasmid consistent with the I179S substitution causing a loss of RelA_P1_ activity (Fig. 3B). It is unclear why the same loss-of-function mutation would occur independently in three lines.

The *relA_P1_* gene is not found in all P1 phages (Figs S2 & 5). In isolates that do encode *relA_P1_* it is found in a variable and comparatively low GC area designated as region of difference 1 (RD-1) that is thought to comprise recently acquired genes (Venturini et al. 2019). To survey the incidence of *relA_P1_* in P1 phages, we used BLASTn to first identify sequence accessions with two characteristic features of P1 phage; the phage replication protein, *repL*, and the P1 plasmid replication protein, *repA* (Johnson et al. 2008; Venturini et al. 2019). Using a cutoff of 90% query coverage and sequence identity, 141 accessions matched both features. Of these, six also matched *relA_P1_*. We also found matches to *relA_P1_* in 25 additional accessions. In all cases, *relA_P1_* matches were flanked by the same *mat* and *lxc* genes found in the M114 P1 element (Figs S2 & 5). These genes are involved in phage related processes—*mat* in phage maturation and *lxc* as a modulator of C1-mediated repression—consistent with most matches occurring in phage or phage-like elements, even if they do not encode *repL* and *repA*. The strong genetic linkage between *relA_P1_* and flanking genes occurring in an otherwise variable region suggests a recent spread of *relA_P1_* among a range of phage.

## Discussion

We investigated the genetic basis of a host-dependent cost of *bla*_TEM-116*\**_ and of compensation to this cost. This ARG was previously demonstrated to impose a significant fitness cost to one *E. coli* host strain, M114, but to have little effect in 11 other tested strains (Lai and Cooper 2023). We found that changes to a P1-like bacteriophage element present in M114, resulting in either loss of the element or loss of function of *relA_P1,_* a gene it encoded, occurred in all of nine evolved clones that compensated the cost of *bla*_TEM-116*\**_. Genetic manipulation experiments demonstrated that the evolved *relA_P1_* mutation was necessary and sufficient to account for compensation of the cost of *bla*_TEM-116*\**_ in M114. Moreover, introduction of *relA_P1_* in a naïve host, REL606, in which *bla*_TEM-116*\**_ was initially neutral, caused *bla*_TEM-116*\**_ to confer a significant cost. These results demonstrate that the cost of *bla*_TEM-116*\**_ seen in this study was the result of an antagonistic interaction between phage encoded *relA_P1_* and *bla*_TEM-116*\**_, highlighting the potential influence of bacterial accessory genomes on the maintenance and evolution of ARGs.

Several mechanisms of β-lactamase cost have been identified. For example, costs of BlaCTX-M-15, BlaSME-1, BlaVIM-1 and BlaSPM-1 depend on their signal peptides which are required to translocate the β-lactamases across the inner membrane to the periplasm and can increase cell envelope stress (Marciano et al. 2007; López et al. 2019; Rajer and Sandegren 2022). Other β-lactamases have low levels of DD-transpeptidase activity, which may impose costs by affecting normal peptidoglycan synthesis (Fernández et al. 2012). We did not determine the mechanistic basis of *bla*_TEM-116*\**_ cost in the M114 strain, but we did demonstrate that its cost depends on the presence of the phage encoded *relA_P1_*. Protein sequence alignments indicate that RelA_P1_ has similarity to the ppGpp synthetase domain of the chromosomal RelA and SpoT enzymes and that it lacks any recognized regulatory domain (Fig. S1). During nutrient starvation *E. coli* ppGpp levels increase sharply via activation of RelA or SpoT ppGpp synthetase activity (Chau et al. 2021). ppGpp regulates various genes involved in cell physiology, resulting in growth arrest as part of a so-called stringent response. In the absence of any regulatory domain, the phage encoded RelA_P1_ may increase the basal level of ppGpp and hypersensitize cells to mild stresses. Hypersensitization could lead to a fitness cost in the presence of BlaTEM-116* if that protein compromised the cell envelope or affected peptidoglycan structure. These changes might be tolerated in the wild type cells but induce the stringent response in RelA_P1_-hypersensitized cells (Roghanian et al. 2019).

The function of *relA_P1_* in the context of the phage P1_M114_ is unclear. A candidate adaptive mechanism for RelA_P1_ functioning as a small ppGpp synthetase enzymes is as a toxin or component of a toxin-antitoxin module (Jimmy et al. 2020; Kurata et al. 2021). For example. a homolog of ppGpp synthase, Tas1, which pyrophosphorylates adenosine nucleotides to produce ppApp, can be delivered via a type VI secretion system to neighboring cells to cause cell death (Ahmad et al. 2019). However, we found that *relA_P1_* expression did not decrease cell fitness, suggesting either that RelA_P1_ is not a toxin or that the hosts already encoded a cognate antitoxin. Alternatively, if RelA_P1_ increases basal ppGpp levels, lowering the threshold for stringent response induction, an encoding phage might gain a selective advantage by early induction of the lytic cycle in harsh environments or through abortive infection providing population-level resistance to secondary infection (Dedrick et al. 2017). Moreover, the elevated basal level of ppGpp may increase the proportion of persister cells induced by stochastic activation of the stringent response (Pacios et al. 2020). Increased spontaneous persister cells may increase the survival of the host population (and hence the temperate phage) in fluctuating environments.

Whatever the basis of the genetic interaction between *relA_P1_* and *bla*_TEM-116*\**_ it is exclusive to that β-lactamase among a set of three that we have considered. In previous work, we found that two other β-lactamases, BlaCTX-M-15 and BlaSHV12, did not confer significant costs when introduced into M114. We note also that one sequenced P1 strain, pTZ20-1P, encodes both *relA_P1_* and *bla*_TEM-1_ (Fig. S2) (Venturini et al. 2019). It would be interesting to investigate if the antagonism between these genes exists in this phage and if not, how it has been resolved. In any case, studying the interaction dynamics of *relA_P1_* and *bla*_TEM-116*\**_ may have practical application in developing approaches to reduce the dissemination of *bla*_TEM-116*\**_ by increasing its cost to host cells.

Models can provide important insight into the implications of strain-specific fitness effects for the ecological dynamics of ARG encoding plasmids. A study that examined the effect of a carbapenemase-carrying plasmid, pOXA-48_K8, over a set of 50 enterobacterial isolated found large differences in its effect on fitness, ranging from a ∼20% benefit to a ∼20% cost across a set of enterobacterial isolates (Valle et al. 2021). Such variation was predicted to be beneficial for the maintenance of the plasmid in a complex bacterial community, allowing it to be maintained in strains in which it confers a low cost in environments not containing the corresponding antibiotic then being transferred and selected in higher-cost strains in environments containing the antibiotic (Valle et al. 2021).

Specific predictions of ARG-plasmid dynamics are, however, complicated by the potential for compensatory mutations to change fitness effects over time (Comas et al. 2012; Harrison et al. 2015; Millan et al. 2015; Loftie-Eaton et al. 2017; Lin et al. 2018; Merker et al. 2018). Even the location—on the host chromosome or on the costly plasmid itself—of compensatory mutations, let alone the effect and rate at which they occur, has been shown to have a major impact on success (Zwanzig et al. 2019). Our demonstration that both the cost of an ARG and the subsequent compensation of that cost can depend on a secondary horizontally mobile element (HME), represents a further complication. If costs and compensation depend on the presence and success of a secondary HME, changes in ARG costs can not only change quickly and independently of host genotype characteristics, but may also be selected unpredictably depending on factors that favour the spread of the HME. Nevertheless, we note that there remain biases in the presence of ARGs in different species – for example, the metallo-β-lactamases BlaVIM-2 and BlaSPM-1 are frequently found in *Pseudomonas aeruginosa* but rarely in other enterobacteria, likely due to differences in their costs – suggesting that not all costs can be overcome (López et al. 2019).

In summary, we demonstrate rapid selection of host cells to compensate for the cost of the *bla*_TEM-116*\**_ resistance determinant. Compensation mutations included deletion or point mutations in a phage encoded gene, *relA_P1_*, which we demonstrate interacts antagonistically with *bla*_TEM-116*\**_. It is unclear if such antagonistic interactions are common, but a survey on the fitness cost of ∼ 200 ARGs has shown that cell-interacting ARGs are more likely to incur a cost (Porse et al. 2018), emphasizing the importance of the compatibility between a HGT gene and its host. Future studies on the identification of the genetic antagonism between an ARG and its host and on the significance of the antagonistic interactions on the dissemination and maintenance of ARGs in bacterial populations will contribute to a comprehensive view on the flow of ARGs between bacteria in nature.

## References

Ahmad S, Wang B, Walker MD, Tran HR, Stogios PJ, Savchenko A, Grant RA, McArthur AG, Laub MT, Whitney JC. 2019. An interbacterial toxin inhibits target cell growth by synthesizing (p)ppApp. Nature 575:674–678.

Andersson D, Hughes D. 2010. Antibiotic resistance and its cost: is it possible to reverse resistance? Nat Rev Micro 8:260–271.

Barrick JE, Kauth MR, Strelioff CC, Lenski RE. 2010. *Escherichia coli rpoB* mutants have increased evolvability in proportion to their fitness defects. Mol Biol Evol 27:1338–1347.

Björkman J, Andersson DI. 2000. The cost of antibiotic resistance from a bacterial perspective. Drug Resist. Updat. 3:237–245.

Chau NYE, Ahmad S, Whitney JC, Coombes BK. 2021. Emerging and divergent roles of pyrophosphorylated nucleotides in bacterial physiology and pathogenesis. PLoS Pathog. 17:e1009532.

Collaborators AR, Murray CJ, Ikuta KS, Sharara F, Swetschinski L, Aguilar GR, Gray A, Han C, Bisignano C, Rao P, et al. 2022. Global burden of bacterial antimicrobial resistance in 2019: a systematic analysis. Lancet Lond Engl 399:629–655.

Comas I, Borrell S, Roetzer A, Rose G, Malla B, Kato-Maeda M, Galagan J, Niemann S, Gagneux S. 2012. Whole-genome sequencing of rifampicin-resistant Mycobacterium tuberculosis strains identifies compensatory mutations in RNA polymerase genes. Nat. Genet. 44:106–110.

Davis BH, Poon AFY, Whitlock MC. 2009. Compensatory mutations are repeatable and clustered within proteins. Proc. R. Soc. B: Biol. Sci. 276:1823–1827.

Deatherage DE, Barrick JE. 2014. Identification of mutations in laboratory-evolved microbes from next-generation sequencing data using breseq. Methods Mol Biology 1151:165 188.

Dedrick RM, Jacobs-Sera D, Bustamante CAG, Garlena RA, Mavrich TN, Pope WH, Reyes JCC, Russell DA, Adair T, Alvey R, et al. 2017. Prophage-mediated defence against viral attack and viral counter-defence. Nat. Microbiol. 2:16251.

Ellis, Haaland P, Hahne F, Meur NL, Gopalakrishnan N, Spidlen J, Jiang M, Finak G. 2022. flowCore: Basic structures for flow cytometry data. R package version 2.8.0.

Fernández A, Pérez A, Ayala JA, Mallo S, Rumbo-Feal S, Tomás M, Poza M, Bou G. 2012. Expression of OXA-type and SFO-1 β-lactamases induces changes in peptidoglycan composition and affects bacterial fitness. Antimicrob Agents Ch 56:1877–1884.

Gallagher RR, Li Z, Lewis AO, Isaacs FJ. 2014. Rapid editing and evolution of bacterial genomes using libraries of synthetic DNA. Nat Protoc 9:2301–2316.

Gelder LD, Ponciano JM, Abdo Z, Joyce P, Forney LJ, Top EM. 2004. Combining mathematical models and statistical methods to understand and predict the dynamics of antibiotic-sensitive mutants in a population of resistant bacteria during experimental evolution. Genetics 168:1131–1144.

Goodman DB, Church GM, Kosuri S. 2013. Causes and effects of N-terminal codon bias in bacterial genes. Science 342:475–479.

Harrison E, Guymer D, Spiers AJ, Paterson S, Brockhurst MA. 2015. Parallel compensatory evolution stabilizes plasmids across the parasitism-mutualism continuum. Curr. Biol. 25:2034–2039.

Jain R, Rivera MC, Lake JA. 1999. Horizontal gene transfer among genomes: The complexity hypothesis. Proc. Natl. Acad. Sci. 96:3801–3806.

Jiang Y, Chen B, Duan C, Sun B, Yang J, Yang S. 2015. Multigene editing in the *Escherichia coli* genome via the CRISPR-Cas9 system. Appl Environ Microb 81:2506–2514.

Jimmy S, Saha CK, Kurata T, Stavropoulos C, Oliveira SRA, Koh A, Cepauskas A, Takada H, Rejman D, Tenson T, et al. 2020. A widespread toxin−antitoxin system exploiting growth control via alarmone signaling. Proc. Natl. Acad. Sci. USA 117:10500–10510.

Johnson M, Zaretskaya I, Raytselis Y, Merezhuk Y, McGinnis S, Madden TL. 2008. NCBI BLAST: a better web interface. Nucleic Acids Res. 36:W5–W9.

Kudla G, Murray AW, Tollervey D, Plotkin JB. 2009. Coding-sequence determinants of gene expression in *Escherichia coli*. Science 324:255–258.

Kurata T, Brodiazhenko T, Oliveira SRA, Roghanian M, Sakaguchi Y, Turnbull KJ, Bulvas O, Takada H, Tamman H, Ainelo A, et al. 2021. RelA-SpoT Homolog toxins pyrophosphorylate the CCA end of tRNA to inhibit protein synthesis. Mol. Cell 81:3160–3170.e9.

Lai H-Y, Cooper TF. 2023. Costs of antibiotic resistance genes depend on host strain and environment and can influence community composition. bioRxiv 2023.09.07.556771

Lin W, Zeng J, Wan K, Lv L, Guo L, Li X, Yu X. 2018. Reduction of the fitness cost of antibiotic resistance caused by chromosomal mutations under poor nutrient conditions. Environ. Int. 120:63–71.

Loftie-Eaton W, Bashford K, Quinn H, Dong K, Millstein J, Hunter S, Thomason MK, Merrikh H, Ponciano JM, Top EM. 2017. Compensatory mutations improve general permissiveness to antibiotic resistance plasmids. Nat Ecol Evol 1:1354–1363.

López C, Ayala JA, Bonomo RA, González LJ, Vila AJ. 2019. Protein determinants of dissemination and host specificity of metallo-β-lactamases. Nat Commun 10:3617.

Marciano DC, Karkouti OY, Palzkill T. 2007. A fitness cost associated with the antibiotic resistance enzyme SME-1 β-lactamase. Genetics 176:2381–2392.

McKenzie GJ, Craig NL. 2006. Fast, easy and efficient: site-specific insertion of transgenes into Enterobacterial chromosomes using Tn*7* without need for selection of the insertion event. Bmc Microbiol 6:39.

Merker M, Barbier M, Cox H, Rasigade J-P, Feuerriegel S, Kohl TA, Diel R, Borrell S, Gagneux S, Nikolayevskyy V, et al. 2018. Compensatory evolution drives multidrug-resistant tuberculosis in Central Asia. eLife 7:e38200.

Millan AS, Toll-Riera M, Qi Q, MacLean RC. 2015. Interactions between horizontally acquired genes create a fitness cost in *Pseudomonas aeruginosa*. Nat. Commun. 6:6845.

Moradigaravand D, Palm M, Farewell A, Mustonen V, Warringer J, Parts L. 2018. Prediction of antibiotic resistance in *Escherichia coli* from large-scale pan-genome data. PLoS Comput. Biol. 14:e1006258.

Nguyen TN, Phan QG, Duong LP, Bertrand KP, Lenski RE. 1989. Effects of carriage and expression of the Tn10 tetracycline-resistance operon on the fitness of *Escherichia coli* K12. Mol Biol Evol 6:213–225.

Nyerges Á, Csörgő B, Nagy I, Bálint B, Bihari P, Lázár V, Apjok G, Umenhoffer K, Bogos B, Pósfai G, et al. 2016. A highly precise and portable genome engineering method allows comparison of mutational effects across bacterial species. Proc National Acad Sci USA 113:2502–2507.

Pacios O, Blasco L, Bleriot I, Fernandez-Garcia L, Ambroa A, López M, Bou G, Cantón R, Garcia-Contreras R, Wood TK, et al. 2020. (p)ppGpp and Its role in bacterial persistence: New challenges. Antimicrob. Agents Chemother. 64:e01283–20.

Park C, Zhang J. 2012. High expression hampers horizontal gene transfer. Genome Biol Evol 4:523–532.

Popa O, Hazkani-Covo E, Landan G, Martin W, Dagan T. 2011. Directed networks reveal genomic barriers and DNA repair bypasses to lateral gene transfer among prokaryotes. Genome Res 21:599–609.

Porse A, Schou TS, Munck C, Ellabaan MMH, Sommer MOA. 2018. Biochemical mechanisms determine the functional compatibility of heterologous genes. Nat. Commun. 9:522.

R Core Team. 2021. R: A language and environment for statistical computing. Available from: https://www.R-project.org/

Rahman S, Kesselheim AS, Hollis A. 2023. Persistence of resistance: a panel data analysis of the effect of antibiotic usage on the prevalence of resistance. J. Antibiot. 76:270–278.

Rajer F, Sandegren L. 2022. The role of antibiotic resistance genes in the fitness cost of multiresistance plasmids. Mbio 13:e03552–21.

Rivera MC, Jain R, Moore JE, Lake JA. 1998. Genomic evidence for two functionally distinct gene classes. Proc. Natl. Acad. Sci. USA 95:6239–6244.

Roghanian M, Semsey S, Løbner-Olesen A, Jalalvand F. 2019. (p)ppGpp-mediated stress response induced by defects in outer membrane biogenesis and ATP production promotes survival in Escherichia coli. Sci. Rep. 9:2934.

Sorek R, Zhu Y, Creevey CJ, Francino MP, Bork P, Rubin EM. 2007. Genome-wide experimental determination of barriers to horizontal gene transfer. Science 318:1449–1452.

Szamecz B, Boross G, Kalapis D, Kovács K, Fekete G, Farkas Z, Lázár V, Hrtyan M, Kemmeren P, Koerkamp MJAG, et al. 2014. The genomic landscape of compensatory evolution. Plos Biol 12:e1001935.

Valle AA, León-Sampedro R, Rodríguez-Beltrán J, DelaFuente J, Hernández-García M, Ruiz-Garbajosa P, Cantón R, Peña-Miller R, Millán AS. 2021. Variability of plasmid fitness effects contributes to plasmid persistence in bacterial communities. Nat Commun 12:2653.

Venturini C, Zingali T, Wyrsch ER, Bowring B, Iredell J, Partridge SR, Djordjevic SP. 2019. Diversity of P1 phage-like elements in multidrug resistant *Escherichia coli*. Sci. Rep. 9:18861.

Wang Y, Arenas CD, Stoebel DM, Flynn K, Knapp E, Dillon MM, Wünsche A, Hatcher PJ, Moore FB-G, Cooper VS, et al. 2016. Benefit of transferred mutations is better predicted by the fitness of recipients than by their ecological or genetic relatedness. Proc National Acad Sci USA 113:5047–5052.

Wein T, Hülter NF, Mizrahi I, Dagan T. 2019. Emergence of plasmid stability under non-selective conditions maintains antibiotic resistance. Nat. Commun. 10:2595.

Zaslaver A, Bren A, Ronen M, Itzkovitz S, Kikoin I, Shavit S, Liebermeister W, Surette MG, Alon U. 2006. A comprehensive library of fluorescent transcriptional reporters for *Escherichia coli*. Nat Methods 3:623–628.

Zwanzig M, Harrison E, Brockhurst MA, Hall JPJ, Berendonk TU, Berger U. 2019. Mobile compensatory mutations promote plasmid survival. Msystems 4: e000186–18.

