## Supplementary Information for "Interactions with a phage gene underlie costs of a β-lactamase"

**Table S1. Oligonucleotides used to introduce mutations into the P1-like phage gene *relA_P1***

| Oligo ID | Purpose | Sequence (5' – 3') |
|---|---|---|
| MAGE_relA_anc_o1 | Revert evolved *relA_P1*$^{I179S}$ allele | C*T*TCGATCAGAAAGAGGATCAATATCCGTGGCGAAAAACAAAGATTGAGATACAGTTAAGAACTCAATTGCAGCATGCTTGGGCTACTAG |
| MAGE_relA_null_o1 | Introduce null *relA_P1* allele (incorporating consecutive stop codons) | C*T*TCGATCAGAAAGAGGATCAATATCCGTGGCGAtAAACAtAGATTtAGATAtAGTTAAGAACTCAATTGCAGCATGCTTGGGCTACTAG |
| MAGE_relA_evo_o1 | Introduce evolved *relA_P1*$^{I179S}$ allele | C*T*TCGATCAGAAAGAGGATCAATATCCGTGGCGAAAAACAAAGAgTGAGATACAGTTAAGAACTCAATTGCAGCATGCTTGGGCTACTAG |

* phosphorothioated nucleotides

Lower-case nucleotides indicate changes relative to the ancestral allele.

Table S2. Mutations identified in evolved plasmid-carrying clones.

| Control (pmFP) vector lines | | | | | | pmFP-*bla*$_{\text{TEM-116*}}$ lines | | | | | | | | | Gene | Description |
|---|---|---|---|---|---|---|---|---|---|---|---|---|---|---|---|---|
| 1 | 2 | 3 | 4 | 5 | 6 | 1 | 2 | 3 | 4 | 5 | 6 | 7 | 8 | 9 | | |
| Δ1bp | Δ72bp | | | Δ1bp | | | | | | | | | | | *GIAEAHOA_00676* | O-antigen polymerase |
| | | | K36* | | | | | | | | | | | | *rfaL* | O_antigen ligase |
| | | P52S | | | | | | | | | | | | | *dut* | dUTPase |
| R562C | E82* | R62L | | R562C | R562C | | | | L92Q | Q280K | | | | +1bp | *prkA* | Serine protein kinase |
| | | | | | | | | | | | | A196V | | | *ydhP* | MFS transporter |
| | | | | | | Q407* | | | | | | | S318P | | *yhbH* | Probable sigma(54) modulation protein |
| | | | | | | | | | | +2bp | | | | | *GIAEAHOA_00573* | Phage tail protein |
| | | | | | | | H210Y | | | | | | | | *GIAEAHOA_02332* | Glycosyltransferase |
| | | | | | | | | | | | | | D58Y | | *nfsA* | Nitroreductase |
| | | | | | | | | | | | D54bp | | | | *ycgB* | Putative SpoVR family protein |
| | | | | | | Loss | Loss | | Loss | Loss | Loss | | | Loss | *Phage P1_M114* | Phage P1_M114 |
| | | | | | | | | I179S | | | | I179S | I179S | | *relAP1* | Putative small ppGpp synthase |

* Intergenic and synonymous mutations are not excluded from this table.

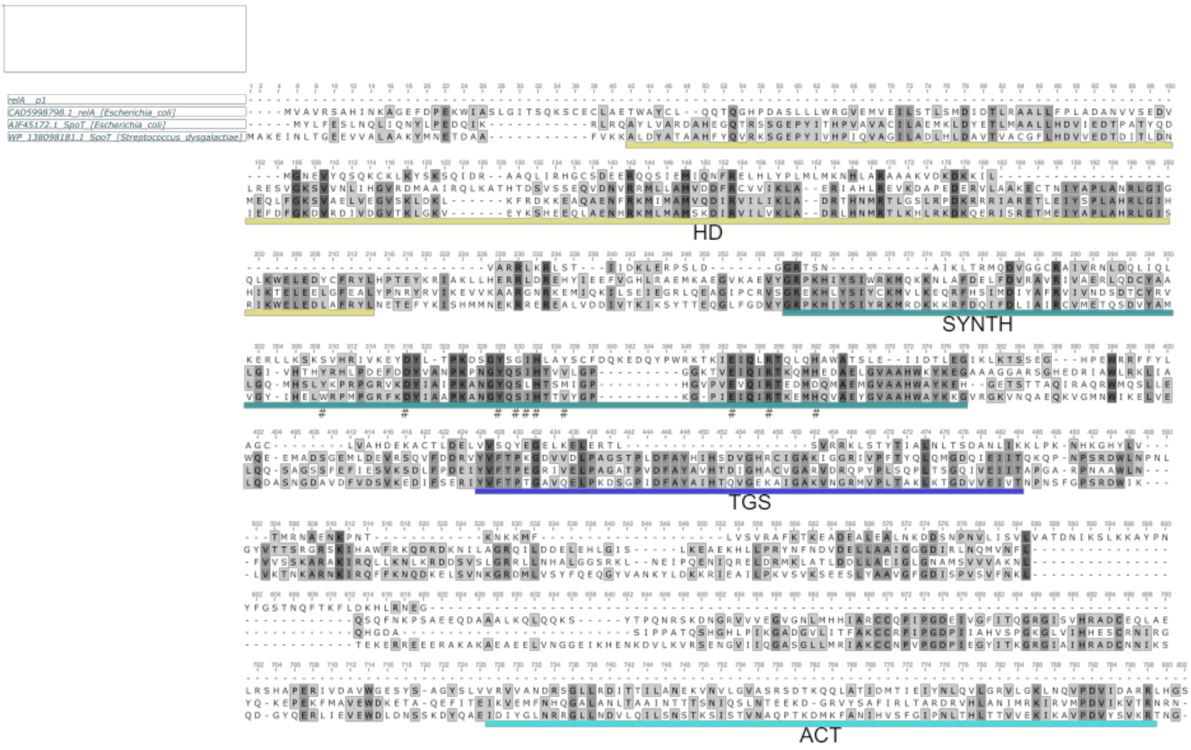


**Figure S1 Protein alignment of RelA/SpoT homologs.** Protein homologs of RelA/SpoT from different species origins were aligned with CLUSTAL W 2.0. SpoT from *Streptococcus dysgalactiae* is a bifunctional protein with both ppGpp synthetase (SYNTH, green line) and hydrolase (HD, yellow line) activity in the N-terminal domain. RelA from *E. coli* only has the synthetase activity, and SpoT from *E. coli* has both synthetase and hydrolase activity. The C-terminal region of the full length RelA/SpoT proteins contains regulatory domains (TGS, blue line, and ACT, cyan line) that binds to other elements such as ribosomes. RelA$_{P1}$ from phage P1 aligns to the N-terminal domain of the full length RelA/SpoT. The RelA$_{P1}$-I179S mutation is at position 352 on this alignment. Amino acids critical for the synthetase activity around I179 of RelA$_{P1}$ are marked with the hash symbol (Hogg et al. 2004; Atkinson et al. 2011).

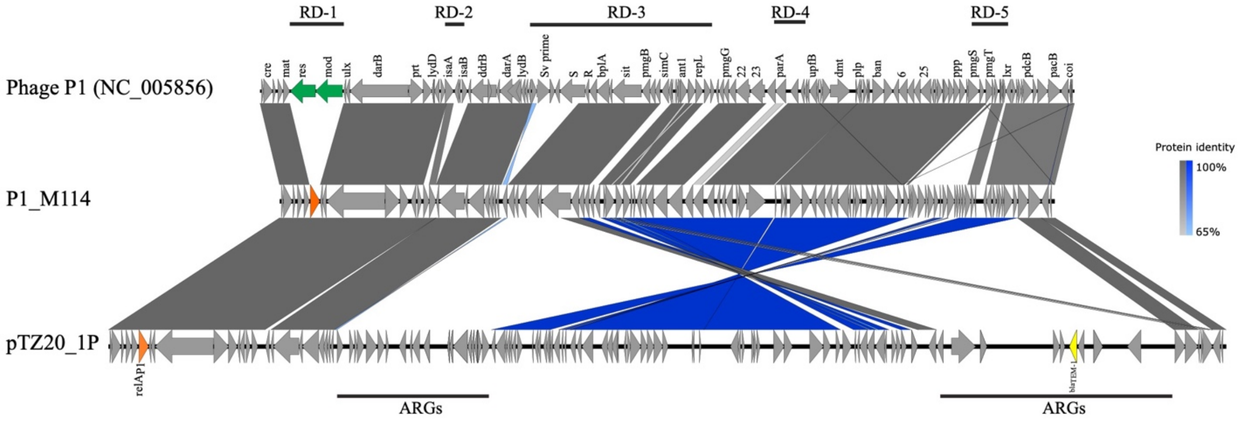


**Figure S2. Genetic map and alignment of phage P1 variants.** Genome comparison between phage P1 variants. Regions noted as RD1-5 indicate areas of sequence difference between the lab strain and *a range of* P1 *isolates* (Venturini et al. 2019). The protein sequence identity between $P1_{M114}$ and pTZ20_1P or the lab strain *(NC_005856)* is colored by grey or blue (inverted region) color gradients. The $relA_{P1}$ gene (orange arrow) is present in P1_M114 and pTZ20_1P but absent in the corresponding region of the lab strain phage (alternative genes are identified with green arrows). pTZ20_1P has acquired multiple ARGs, including $\textit{bla}_{TEM\text{-}1}$ (yellow arrow).

### References


Atkinson GC, Tenson T, Hauryliuk V. 2011. The RelA/SpoT homolog (RSH) superfamily: Distribution and functional evolution of ppGpp synthetases and hydrolases across the tree of life. *PLoS ONE* 6:e23479.

Hogg T, Mechold U, Malke H, Cashel M, Hilgenfeld R. 2004. Conformational antagonism between opposing active sites in a bifunctional RelA/SpoT homolog modulates (p)ppGpp metabolism during the stringent response. *Cell* 117:57–68.

Venturini C, Zingali T, Wyrsch ER, Bowring B, Iredell J, Partridge SR, Djordjevic SP. 2019. Diversity of P1 phage-like elements in multidrug resistant Escherichia coli. *Sci. Rep.* 9:18861.
